# Unexpected delayed incursion of highly pathogenic avian influenza H5N1 (clade 2.3.4.4b) in the Antarctic region

**DOI:** 10.1101/2023.10.24.563692

**Authors:** Simeon Lisovski, Anne Günther, Meagan Dewar, David Ainley, Fabián Aldunate, Rodrigo Arce, Grant Ballard, Silke Bauer, Josabel Belliure, Ashley C. Banyard, Thierry Boulinier, Ashley Bennison, Christina Braun, Craig Cary, Paulo Catry, Augustin Clessin, Maelle Connan, Edna Correia, Aidan Cox, Juan Cristina, Megan Elrod, Julia Emerit, Irene Ferreiro, Zoe Fowler, Amandine Gamble, José P. Granadeiro, Joaquin Hurtado, Dennis Jongsomjit, Célia Lesage, Mathilde Lejeune, Amanda Kuepfer, Amélie Lescroël, Amy Li, Ian R. McDonald, Javier Menéndez-Blázquez, Virginia Morandini, Gonzalo Moratorio, Teresa Militão, Pilar Moreno, Paula Perbolianachis, Jean Pennycook, Maryam Raslan, Scott M. Reid, Roanna Richards-Babbage, Annie E. Schmidt, Martha Maria Sander, Lucy Smyth, Alvaro Soutullo, Andrew Stanworth, Léo Streith, Jérémy Tornos, Arvind Varsani, Ulrike Herzschuh, Martin Beer, Michelle Wille

## Abstract

The current highly pathogenic avian influenza H5N1 panzootic has substantial impacts on wild birds and marine mammals. Although major outbreaks occurred in South America, incursion to Antarctica emerged late in the breeding season of 2023/2024 and was confined the wider region of the Antarctic Peninsula. To infer potential underlying processes, we compiled H5N1 surveillance from Antarctica and Sub-Antarctic Islands prior to the first confirmed cases.

## Main text

The increasing intensity of highly pathogenic avian influenza virus (HPAIV) H5N1 clade 2.3.4.4b outbreaks have had a substantial impact on poultry and wildlife ^1^. Wild bird movements have underpinned the rapid spread of this virus that swept across most continents, except for Australia and Antarctica, within two years ^2^. Compared to previous HPAIV subtypes and clades, H5N1 2.3.4.4b has significantly improved replication in wild birds ^3^, and increased fitness through continuous reassortments ^4^ which has likely contributed to a shift in infection dynamics leading to the infection of a broader range of avian species ^1^. In addition to their role as viral spreaders, wild birds are suffering huge losses following mass mortality events, and the scale of mortality amongst wild birds is likely in the millions rather than tens of thousands reported ^5^. Thus, the recent panzootic is a serious conservation concern for a large range of wild bird species.

Due to the absence of waterfowl species that migrate to the Antarctic and sub-Antarctic islands, the incursion risk of HPAIV in these southernmost regions had been considered low prior to 2021. However, waterfowl are present in northern fringes of the Southern Ocean, and millions of known migration and post-breeding dispersal routes establish links and thereby substantial global connectivity, including with regions of recent HPAI H5N1 outbreaks involving seabirds and marine mammals ^2^. Despite the perceived remoteness, low pathogenicity avian influenza viruses and antibodies against these viruses have previously been detected in various seabird species nesting at sites along the Antarctic Peninsula and South Shetland Islands, with viral genomes illustrating phylogenetic connectivity to viruses circulating on other continents ^6,7^. As a result, the Scientific Committee on Antarctic Research (SCAR) Antarctic Wildlife Health Network (AWHN) had considered the risk of incursion of the recent panzootic HPAIV H5 into the Antarctic region in 2022/23 summer season to be high ^8^, and considerably higher in 2023/24 following virus spread to the southernmost regions of South America ^9^, infecting seabirds including Magellanic penguins (*Spheniscus magellanicus*) and Humbold penguins (*Spheniscus humboldti*), and several species of marine mammals (9).

To identify possible incursions of H5N1 into the Antarctic region during the summer season 2022/23 and the early season 2023/24, we sampled migratory seabirds at different locations across Antarctica and in sub-Antarctic areas (Figure 1), and collated a range of observation data. Here, we define Antarctica as the region south of the Antarctic Convergence and sub-Antarctic areas include adjacent Islands within the Southern Ocean. In particular, we aimed to collect information pertaining to suspicious signs of unusual mortality and known clinical signs of HPAIV infection including loss of coordination and balance, trembling head and body, lethargy, respiratory distress, and conjunctivitis ^8^. Across all locations, sample collection was done in accordance with institutional animal ethics approval and sample testing was performed with national frameworks, with details available in the technical annex.

**Figure 1:**
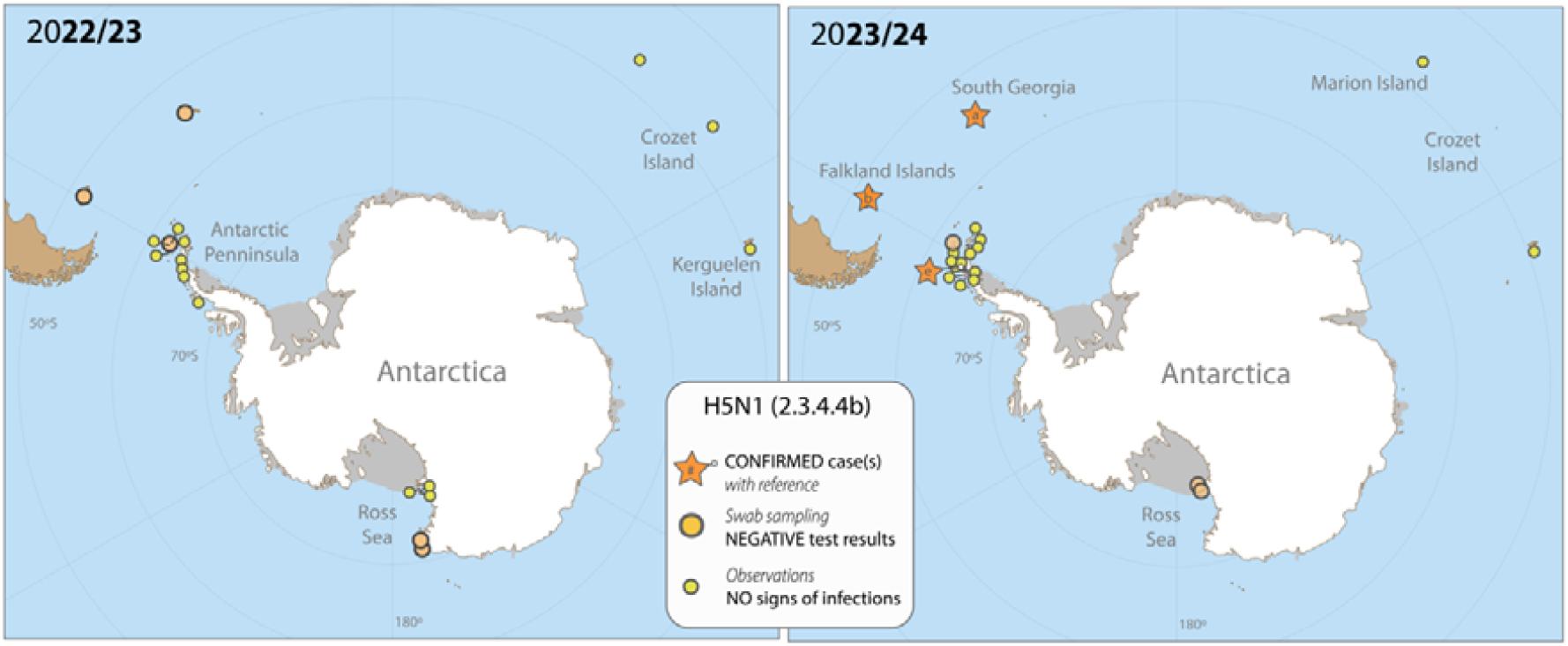
Sampling locations for qPCR analysis and the detection of H5N1 2.3.4.4b, as well as locations with intensive observational efforts to identify signs of HPAI infections within breeding bird communities for the breeding season 2022/23 (left) and 2023/24 (right). In addition, locations of confirmed cases of infection in 2023/24 (left) are included. Numbers refer to the following references, (a) technical annex, (b) Bennison et al. ^10^, (c) First confirmed case near Argentina’s Primavera research station ^11,12^. Maps created with Natural Earth.

Overall, sampling and observational efforts were conducted from early November 2022 to late March 2023, and from October 2023 onwards until the end of February in 2024. Surveillance efforts included a large range of species (*i.e*., penguins, gulls, skuas, and petrels; see technical annex for more information) and locations. In 2022/23, samples for HPAIV testing were collected from apparently healthy birds from 20 locations in the sub-Antarctic and Antarctic Region. There were several suspicious observations of dead wild birds on the Falkland Islands (Gentoo penguin *Pygoscelis papua*, Cattle egret *Bubulcus ibis*), and South Georgia (Wandering albatrosses *Diomedea exulans*). However, all swab samples collected from these animals, in addition to apparently healthy wild birds in other locations were negative for HPAI (see technical annex for details on location and species). Together, this strongly suggests that HPAIV H5N1 clade 2.3.4.4b did not enter the Antarctic region during the austral summer 2022/23, and that the lack of detection was unlikely due to lack of surveillance, testing or disease investigations. This is in contrast to the seabird breeding season 2023/24. In October 2023 the first confirmed H5N1 cases were detected on the Falkland (Malvinas) Islands, and in November on South Georgia Island in the sub-Antarctic ^10,11^ (Figure 1). Given the overlap of species breeding and migrating via the Falkland Island and South Georgia towards the Antarctic Peninsula and its offshore Islands (e.g., the South Shetland Islands), researchers in the region and the tourist industry have been very diligent in identifying unusual bird behaviour and mortality events. Despite active cases in the Falkland Islands and South Georgia Island, sample collection and observations from 16 locations between November 2023 – early February 2024 in the Antarctic Peninsula and related island were negative for HPAIV. Data from the SCAR monitoring project did, however, report suspected cases in the Antarctic region starting in December 2023 ^11^. These include Brown Skuas on the South Orkney Islands in December 2023 (no samples collected), a mortality in Brown Skuas on Heronia Island in December 2023 (samples collected, HPAIV negative). Since mid-February first positive cases were reported from the Antarctic Peninsula (see Figure 1) ^11,12^. This suggests that H5N1 was spread among colonies in the later breeding season, however, so far there is no evidence for large outbreaks on the Antarctic Peninsula. Further, based on observation data, the strain did not appear to have reached the Indian Ocean sub-Antarctic islands as of February 2024 (see technical annex).

Obviously, incursion risk, and successful establishment of HPAIV is contingent on a combination of factors. Most importantly, that (i) host species get in contact with HPAIV before travelling into the Antarctic regions, (ii) can migrate with an infection, and (iii) have contact and transmit the virus to susceptible species which could be the starting point of a new epizootic. Most species occupying the Antarctic region are pelagic seabirds with little to no contact with terrestrial birds such as waterfowl, significantly reducing their exposure to outbreaks on land (e.g. South America). However, some species like the Brown skua (*Stercorarius antarcticus*) and the giant petrel species (*Macronectes sp.)* are known scavengers (Figure 2), leading to high risks of exposure to HPAIV via the consumption of infected carcasses. It is thus no surprise that Brown skuas where often among the first confirmed cases both on South Georgia and the Antarctic Peninsula ^10^. This species, which can be observed at shorelines of South America, the Falkland Islands and South Georgia ^13^, is likely to be an important player in spreading the virus. Yet, it seems that the connectivity established by the animals’ movements from South America and South Georgia over the Drake Passage to Antarctica is rather limited during the breeding season but might increase again towards the end when the breeding activities terminate, and the movement range of both adults and first juveniles are becoming larger again. Together with the increasing number of naïve juveniles and concomitant changes in densities, this may explain the delay between initial outbreaks in the Falklands/South Georgia and the first confirmed cases on the Antarctic peninsula.

**Figure 2:**
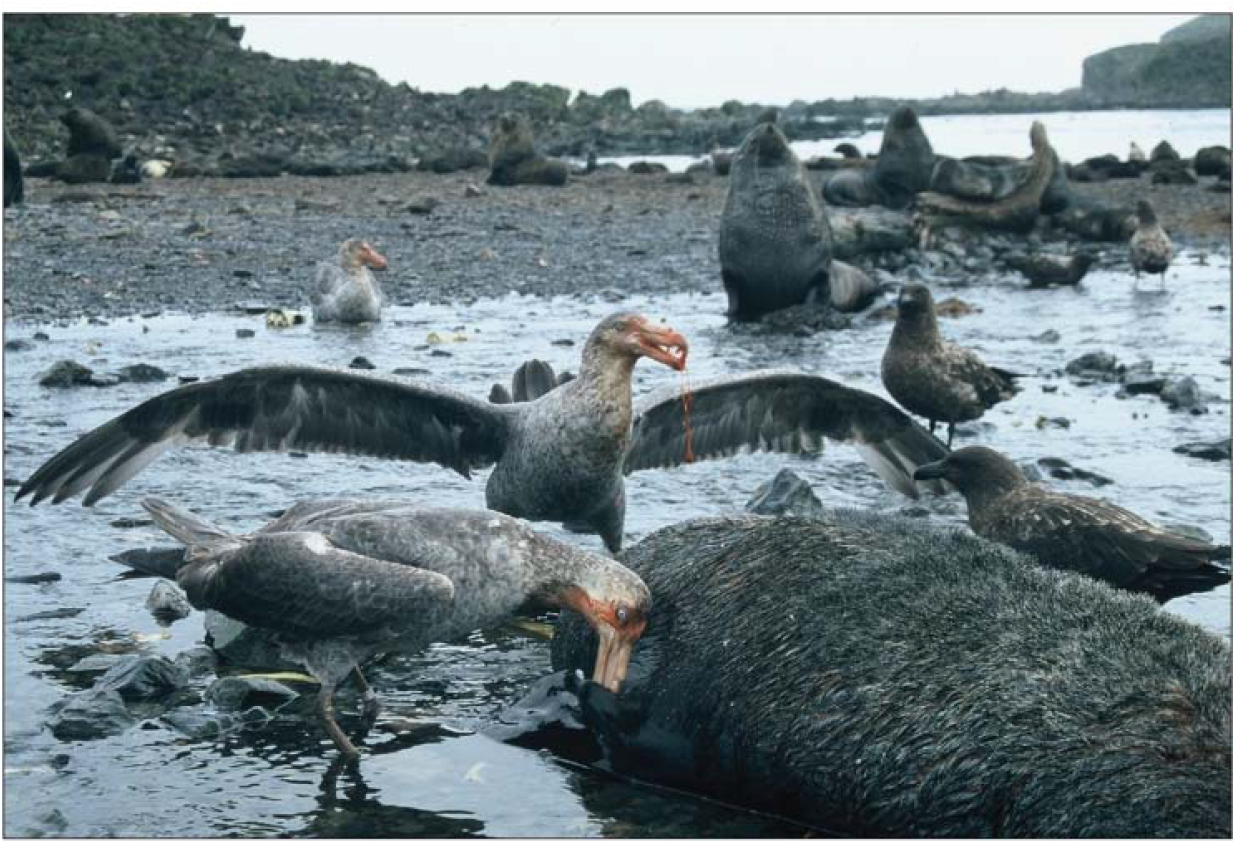
Northern giant petrels and Brown skuas scavenging on an Antarctic fur seal carcass, showing inter-species interactions with the potential for HPAI virus transmission (photo taken on South Georgia by Paulo Catry).

Still, the consequences of viral incursion(s) into Southern Ocean wildlife are unclear but based on observations from other regions, will likely have devastating effects. Critically, densities of seabird colonies are very high, facilitating the transmissions between individuals ^14^. Further prospecting movements of potential recruits, predator-prey interactions between bird species (e.g., skuas, penguins, and sheathbills), as well as species scavenging on dead seabirds and mammals, may promote rapid spread of the virus between colonies ^15^. Once the virus has been established in the region, interaction between seabirds and marine mammals may also result in further transmissions and facilitate the adaptation of the virus in mammalian species ^14^. Finally, most animals of the Southern Ocean are endemic to the region, such that mass mortality events in Antarctica due to HPAIV H5 will cause a very real conservation concern for many species.

Detecting H5N1 incursion(s) and describing the infection dynamics into and within the sub-Antarctic and Antarctic regions is highly relevant and standardized surveys for mortality and sampling should therefore be prioritized. These activities should be undertaken with consideration of the potentially zoonotic risks of (emerging) HPAIV H5 ^8^ and require strict hygiene measures to prevent the spread of the virus through human activities. Sampling and detailed analysis of lineages and virus phenotype will provide crucial information needed to assess risks and respond to future wild bird outbreaks.

## Supporting information

Technical Annex

## Acknowledgements

We thank the following institutions and agencies for financial and logistical support: The Uruguayan Antarctic Institute; Government South Georgia and South Sandwich Islands; Antarctica New Zealand; French Polar Institute (ECOPATH-1151); National Nature Reserve of Terres Australes et Antarctique Françaises; The South African Departments of Science and Innovation (National Research Foundation - South African National Antarctic Programme project no. 129226; MC) and of Forestry, Fisheries and the Environment; Falklands Conservation, the Wildlife Conservation Society; Falkland Islands Government; The Antarctic Research Trust; The South Atlantic Environmental Research Institute; Commission for the Conservation of Antarctic Marine Living Resources (CCAMLR); Spanish Research Agency (VM). The study was supported by the following research grants: ANR (ECOPATHS ANR-21-CE35-0016 and REMOVE_DISEASE ANR-21-BIRE-006-01; TB, ML, JT); Biodiversa and Water JPI under the BiodivRestore ERA-NET Cofund (GA N°101003777; TB, MC, ML, AC, PC); National Science Foundation award ANT 1935870 and ANT 2040199 (GB, DA, ME, AC AS, AL, AL, AV, JP); Royal Society (Newton International Fellowship NIF\R1\211869; AGa); Biodiversity Challenge Fund (Darwin Plus Grant DPLUS167; AGa); Fundação para a Ciência e a Tecnologia (FTC) – Portugal (UIDB/04292/2020 (MARE) and LA/P/0069/2020 (ARNET) and DivRestore/0012/2020 (REMOVE_DISEASE project), Agencia Nacional de Investigación e Innovación’s (ANII) Clemente Estable Fund (project FCE_1_2021_1_166587); Ecos-Sud Program (project PU20B01/U20B03); General Capacity Building Fund; the Department for Environment, Food and Rural Affairs (Defra, UK) and the devolved administrations of Scotland and Wales under grant numbers SV3006, SE2213 and SV3045; Biotechnology and Biological Sciences Research Council (BBSRC) and Defra funded research initiative ‘FluMAP’ [grant number BB/X006204/1, ACB, SMR].

European Union Horizon Europe research program (grant agreement “Kappa-Flu No. 101084171”, with the Schweizer Staatssekretariat, AGü, MB, SB). In addition, we thank Corisande Abiven, Katharina Reusch, Louise Pole-Evans, Edna Correia, Jose P Granadeiro, Michelle Risi and Christopher Jones for fieldwork support, and all the inhabitants and visitors of the Falkland Islands for reporting suspected cases and facilitating research on their lands, in particular Kicki Ericson and Thies Matzen, Sarah Crofts and Micky Reeves, and the Rendell, Pole-Evans, Gould, Delignieres and Hazel families.

## Notes

### Competing Interest Statement

The authors have declared no competing interest.

### Summary of Updates

Correction of references and figure update

