## Supplementary material for "Unexpected delayed incursion of highly pathogenic avian influenza H5N1 (clade 2.3.4.4b) in the Antarctic region": Technical Annex

Observations

Following the guideline of Dewar et al. ^1^, observations on the breeding sites of seabirds in Antarctica and on sub-Antarctic islands included the following signs of potential HPAI infections:

- Neurological issues such as loss of coordination and balance,
- Trembling head and body
- Sudden and rapid increase in the number of birds found dead between
- visits,
- Lethargy and depression, unresponsiveness, lying down, drooping wings, dragging legs,
- Swollen head,
- Closed and excessively watery eyes, possibly with opaque cornea or darkened iris (new clinical signs associated with current outbreak) ^2^,
- Twisting of the head and neck,
- Haemorrhages on shanks of the legs and under the skin of the neck,
- Respiratory distress such as gaping (mouth breathing), nasal snicking (coughing sound), sneezing, gurgling, or rattling, and
- Discoloured or loose watery droppings, bright green in some species.

Observations where mainly done including the whole range of breeding seabird species at the sites. Testing life birds was restricted to species available and possible to catch in larger numbers.

Field and laboratory protocols

1. Falkland Islands

Location: 59.30 West, 51.81 South

**Breeding Season 2023/24**

| **Fieldwork** | | | |
| --- | --- | --- | --- |
| Investigators | Period | Type | Species (sample size) |
| Zoe Fowler,  Amanda Kuepfer,  Julia Emerit,  Léo Streith,  Andy Stanworth | Continuously | Opportunistic observation | Observations are done around the islands by inhabitants and visitors, including all breeding birds (and poultry).  **> 80 SUSPECTED CASES** reported and > 50 set of swabs collected and tested from all over the archipelago (including the sites listed below), **> 10 OUTBREAKS OR ISOLATED CASES CONFIRMED** in Southern Fulmars (vagrant), Black browed albatrosses, Brown skuas, Southern rockhopper penguins, Gentoo penguins. |
| Amandine Gamble  Acknowledgement:  Kicki Ericson,  Thies Matzen | 10.10.2023 – 15.10.2024 | Systematic observation | **NEGATIVE**: On West Point (51.35 West, 60.68 South): Black browed albatross, Southern rockhopper penguin, Gentoo penguin, Magellanic penguin, Striated caracara (Brown skua and Imperial shag not on the island yet) |
| Amandine Gamble, Julia Emerit  Edna Correia,  Jose P. Granadeiro  Acknowledgement:  Peter Hover | 15.10.2023 – 04.03.2024 | Systematic observation  Swab sampling: Oropharyngeal & Cloacal Swabs | On New Island (51.71 West, 61.31 South): Black browed albatross, Brown skua, Southern rockhopper penguin, Gentoo penguin, Magellanic penguin, Imperial shag, Striated caracara  Single observations of dead Black browed albatrosses, Brown skuas, Gentoo penguins (swabs taken) |
| Léo Streith | 09.11.2023 – 19.11.2023 | Systematic observation | On Hummock (51.62 West, 60.44 South): Brown skua, Southern rockhopper penguin, Magellanic penguin, Imperial shag, Striated caracara |
| Julia Emerit, Amanda Kuepfer,  Acknowledgement: Andy Stanworth | 22.11.2023 – 05.03.2024 | Systematic observation  Swab sampling: Oropharyngeal & Cloacal Swabs | On Steeple Jason (51.04 West, 61.21 South): Black browed albatross, Brown skua, Southern rockhopper penguin, Gentoo penguin, Magellanic penguin, Imperial shag, Striated caracara  11.23 unusually high number of dead adult Black browed albatrosses (swabs taken), 01.24 continuing high numbers of dead adults and juveniles of Black browed albatrosses, Southern rockhopper penguins and Brown skuas (swabs taken), 03.24 continuing high numbers of dead adults and juveniles of Black browed albatrosses, Southern rockhopper penguins and Gentoo penguins (swabs taken). |
| Julia Emerit | 27.11.2023 – 09.03.2024 | Systematic observation  Swab sampling: Oropharyngeal & Cloacal Swabs | On Bleaker Island (52.22 West, 58.88 South): Brown skua, Southern rockhopper penguin, Gentoo penguin, Magellanic penguin, Imperial shag, Striated caracara  Single observations of dead Gentoo penguins (swabs taken) and one symptomatic Magellanic penguin |
| Amandine Gamble, Julia Emerit | 08.12.2023 – 02.02.2024 | Systematic observation  Swab sampling: Oropharyngeal & Cloacal Swabs | On Saunders Island (51.36 West, 60.08 South): Black browed albatross, Brown skua, Southern rockhopper penguin, Gentoo penguin, Magellanic penguin, Imperial shag, Striated caracara  Single observations of dead Black browed albatrosses and Brown skuas (swabs taken) |
| Julia Emerit | 04.01.2024 – 09.01.2024 | Systematic observation  Swab sampling: Oropharyngeal & Cloacal Swabs | On Pebble Island (51.31 West, 59.60 South): Brown skua, Southern rockhopper penguin, Gentoo penguin, Magellanic penguin, Imperial shag, Striated caracara  Single observations of dead Southern fulmars (vagrant) and Gentoo penguins (swabs taken) |
| Julia Emerit | 11.01.2024 – 15.01.2024 | Systematic observation  Swab sampling: Oropharyngeal & Cloacal Swabs | On Sealion Island (52.43 West, 59.06 South): Brown skua, Southern rockhopper penguin, Gentoo penguin, Magellanic penguin, Imperial shag, Striated caracara  Single observations of dead and symptomatic Gentoo penguins (swabs taken) |
| Julia Emerit | 07.02.2024 – 12.02.2024 | Systematic observation | On Dunbar (51.40 West, 60.56 South): Black browed albatross, Brown skua, Southern rockhopper penguin, Gentoo penguin, Magellanic penguin, Imperial shag, Striated caracara |

| **Laboratory analyses** | | | |
| --- | --- | --- | --- |
| Investigators | Laboratory | Method | Protocol |
| Zoe Fowler,  Acknowledgement:  Kim Finlayson | KEMH Pathology and Food, Water & Environmental Laboratory  St Mary’s Walk, Stanley, Falkland Islands, FIQQ 1ZZ | qPCR | QIAamp Viral RNA Mini Kit using QIACube extraction robot and standard manufacturer’s instructions.    Nucleic extracts screened by real-time RT-PCR using Primer Design Genesig Bird flu kit (catalogue no: Path-H5N1), using the QuantStudio 5 real time PCR platform.  From the kit insert: The Primerdesign genesig Kit for Influenza A virus subtype H5N1 (avian influenza) (H5N1) genomes are designed for the in vitro quantification of H5N1 genomes. The kit is designed to have a broad detection profile. Specifically, the primers represent 100% homology, with over 95% of the NCBI database reference sequences available at the time of design.  The dynamics of genetic variation mean that new sequence information may become available after the initial design. Primerdesign periodically reviews the detection profiles of our kits and, when required, releases new versions. The primers have 100% homology with over 95% of avian H5N1 isolates globally that have been entered into the influenza sequence database in the NCBI database since 2001. The primers have low sequence homology to other influenza subtypes, and the quantification of both subtyping genes ensures an accurate determination of the H5N1 genotype in a single experiment. However, due to the inherent instability of RNA viral genomes, novel emerging sequences may not be detected. |

**Breeding Season 2022/23**

| **Fieldwork** | | | |
| --- | --- | --- | --- |
| Investigators | Period | Type | Species (sample size) |
| Zoe Fowler,  Tom Hart,  Amandine Gamble, Amanda Kuepfer,  Léo Streith | Continuously | Opportunistic observation | **NEGATIVE**: Observations are done around the islands by inhabitants and visitors, including all breeding birds (and poultry). |
| Zoe Fowler |  | Swab sampling: Oropharyngeal & Cloacal Swabs | **NEGATIVE**: 31.01.2024 (approx. 30) Gentoo penguin chicks (Bertha's Beach) in weak state with visible skin lesions (swabs taken), 27.01.2023 >100 dead juvenile gentoos (swabs taken), 13.02.2023 122 dead Gentoo penguin chicks and more sick individuals (swabs taken). 27.04.2023 dead cattle egret (swabs taken), 01.05.2023 sick cattle egret (swabs taken). |
| Amandine Gamble, Augustin Clessin | 15.10.2022 – 10.03.2023 | Systematic observation | **NEGATIVE**: On New Island (51.71 West, 61.31 South): Black browed albatross, Brown skua, Southern rockhopper penguin, Gentoo penguin, Magellanic penguin, Imperial shag, Striated caracara |
| Amandine Gamble, Paulo Catry | 11.12.2022 – 19.12.2022 | Systematic observation | **NEGATIVE**: On Grand Jason (51.06 West, 61.10 South): Black browed albatross, Brown skua, Southern rockhopper penguin, Gentoo penguin, Magellanic penguin, Imperial shag, Striated caracara |
| Amandine Gamble, Paulo Catry,  Amanda Kuepfer | 19.12.2022 – 30.12.2022 | Systematic observation | **NEGATIVE**: On Steeple Jason (51.04 West, 61.21 South): Black browed albatross, Brown skua, Southern rockhopper penguin, Gentoo penguin, Magellanic penguin, Imperial shag, Striated caracara |

| **Laboratory analyses** | | | |
| --- | --- | --- | --- |
| Investigators | Laboratory | Method | Protocol |
| Zoe Fowler  Acknowledgement:  Kim Finlayson | KEMH Pathology and Food, Water & Environmental Laboratory  St Mary’s Walk, Stanley, Falkland Islands, FIQQ 1ZZ | qPCR | See protocol for season 2023/24 |

### 2. Bird Island, South Georgia

Location: 38.06 West, 54.01 South

**Breeding Season 2022/23**

| **Fieldwork** | | | |
| --- | --- | --- | --- |
| Investigators | Period | Type | Species (sample size) |
| Ashley Bennison | 19.12.2022 – 29.12.2022 | Swab sampling: Oropharyngeal & Cloacal Swabs | **NEGATIVE**: Wandering albatross (4); 19/12/2022 Adult wandering albatross in good body condition found dead, no obvious signs of disease or external trauma. 29/12/2022 Three further wandering albatrosses were found dead, including two old chicks close to fledging and one adult. One chick thought to be starved, the other seemingly healthy. Shortly before death, the adult had been observed with the following symptoms: lethargy, uncoordinated movement, drooping of head. Bird Island HPAI response increased to level 2 to signify potential outbreak. Oropharyngeal and cloacal swabs taken from all three birds. |
| Ashley Bennison | 11.2022 – 02.2023 | Opportunistic observations | **NEGATIVE**: Seabird colonies of Bird Island |

| **Laboratory analyses** | | | |
| --- | --- | --- | --- |
| Investigators | Laboratory | Method | Protocol |
| Ashley C. Banyard, Scott M. Reid | Animal and Plant Health Agency, UK | qPCR | Initial M gene screening and then subtyping qRT-PCR assays for HA and NA were undertaken as described here: FluGlobalNet - Protocols (vla.gov.uk):  https://science.vla.gov.uk/fluglobalnet/protocols.html |

### 3. Fildes Peninsula and Ardley Island, King George Island, South Shetland Islands

Location: 58.96 West, 62.21 South

**Breeding Season 2023/24**

| **Fieldwork** | | | |
| --- | --- | --- | --- |
| Investigators | Period | Type | Species (sample size) |
| Simeon Lisovski, Martha M. Sander, Alvaro Soutullo,  Javier Menéndez-Blázquez,  Maryam Raslan | 17.11.2023 – 19.12.2023 | Swab sampling: Oral- & Cloacal Swab (RNALater) | **NEGATIVE:** South polar skua (37), Brown skua (28), Southern giant petrel (33), Chinstrap penguin (15), Gentoo penguin (43), Adélie penguin (9), Antarctic tern (2), Blue-eyed cormorant (4) |
| Simeon Lisovski, Martha M. Sander, Alvaro Soutullo, Christina Braun,  Javier Menéndez-Blázquez,  Maryam Raslan | 17.11.2023 – 22.02.2024 | Opportunistic observation | **NEGATIVE:** Entire seabird breeding community on Fildes Peninsula and Ardley Island |
| Juan Cristina,  Irene Ferreiro,  Joaquin Hurtado,  Gonzalo Moratorio,  Pilar Moreno | 29.01.2024 – 09.02.2024 | Fecal sampling | **NEGATIVE:** Gentoo penguin (30), Chinstrap penguin (2), Brown skua (2) |

| **Laboratory analyses** | | | |
| --- | --- | --- | --- |
| Investigators | Laboratory | Method | Protocol |
| Anne Günther,  Martin Beer | Friedrich-Loeffler-Institut, Isle of Riems, Germany | qPCR | Swab samples have been stored in PrimeStore® MTM by Longhorn Vaccines & Diagnostics. The supernatant was applied for RNA extraction via the NucleoMag® VET Kit by Macherey-Nagel following the manufacturer’s instructions. A heterologous control RNA ^3^ in 1:10 ratio to the sample RNA elution volume has been included. Subsequent generic RT-qPCR screenings targeted the matrix gene of influenza A viruses and were conducted as described by Koethe et al. ^4^. |
| Juan Cristina,  Irene Ferreiro,  Joaquin Hurtado,  Gonzalo Moratorio,  Pilar Moreno | Laboratorio de Virología Molecular, Centro de Investigaciones Nucleares, Facultad de Ciencias, Universidad de la República, Igua 4225, Montevideo 11400, Uruguay | PCR | Each fecal sample was resuspended in 1× phosphate buffered saline (PBS) to obtain a 10% dilution. This suspension was clarified by centrifugation at 6000 × g for 10 min, and 300 𝜇L of the supernatant was used for total ARN extraction using the QIAamp Viral RNA Mini Kit (QIAamp, USA) according to the manufacturer's instructions. Retrotranscription of RNA samples was performed using SuperScript II Reverse Transcriptase (Invitrogen, Life Technologies, USA) according to the manufacturer's recommended protocol. 100 ng of random hexamer primers (Thermo Scientific, USA) were used to allow the cDNA to be used for the detection of different AIV genes and internal control genes.  A specific PCR for a 330 pb region of the nucleoprotein (NP) gene was performed to determine the presence of AIV in each sample. Q5 High-Fidelity 2X Master Mix (New England Biolabs, USA) was used according to the manufacturer's instructions, with the following cycling conditions: an initial denaturation of 98°C for 30 seconds, followed by 35 cycles consisting of denaturation at 98°C for 10 seconds, annealing at 58°C for 30 seconds and extension at 72°C for 20 seconds, and a final extension at 72°C for 5 minutes, followed by a 10°C hold. The primers used in this PCR (NP1200 and NP1529) were described by Lee et al. ^5^. These primers were designed to detect all AIV haemagglutinin (HA) subtypes. |

**Breeding Season 2022/23**

| **Fieldwork** | | | |
| --- | --- | --- | --- |
| Investigators | Period | Type | Species (sample size) |
| Simeon Lisovski,  Ulrike Herzschuh | 12.11.2022 – 29.11.2022 | Swab sampling: Oral- & Cloacal Swab (RNALater) | **NEGATIVE**: South polar skua (30), Brown skua (30), Southern giant petrel (15), Chinstrap penguin (25), Gentoo penguin (25), Adélie penguin (25) |
| Simeon Lisovski,  Ulrike Herzschuh,  Alvaro Soutullo | 12.11.2022 – 15.01.2023 | Opportunistic observation | **NEGATIVE**: Entire seabird breeding community on Fildes Peninsula and Ardley Island |
| Rodrigo Arce,  Juan Cristina,  Irene Ferreiro,  Joaquin Hurtado, Gonzalo Moratorio, P. Perbolianachis, Pilar Moreno | 30.01.2023 – 12.02.2023 | Fecal sampling | **NEGATIVE**: Gentoo penguin (32), Chinstrap penguin (7), Emperor penguin (1), Brown skua (3) |

| **Laboratory analyses** | | | |
| --- | --- | --- | --- |
| Investigators | Laboratory | Method | Protocol |
| Anne Günther,  Martin Beer | Friedrich-Loeffler-Institut, Isle of Riems, Germany | qPCR | Swab samples have been stored in RNAlater. The supernatant was applied for RNA extraction via the QIAamp Viral RNA Mini Kit by Qiagen following the manufacturer's instructions. For preceding steps see protocol for season 2023/24. |
| Rodrigo Arce,  Juan Cristina,  Irene Ferreiro,  Joaquin Hurtado, Gonzalo Moratorio, Paula Perbolianachis, Pilar Moreno | Laboratorio de Virología Molecular, Universidad de la República, Igua 4225, Montevideo 11400, Uruguay | qPCR | See protocol for season 2023/24 |

### 4. Avian Island

Location: 68.88 West, 67.77 South

**Breeding Season 2022/23**

| **Fieldwork** | | | |
| --- | --- | --- | --- |
| Investigators | Period | Type | Species (sample size) |
| Virginia Morandini,  Josabel Belliure | 23.01.2023 | Observation | **NEGATIVE**: Adelie penguin, entire breeding colony |

### 5. Cape Adare, Victoria Land

Location: 170.19 East, 71.35 South

**Breeding Season 2022/23**

| **Fieldwork** | | | |
| --- | --- | --- | --- |
| Investigators | Period | Type | Species (sample size) |
| Craig Cary | 01.11.2022 – 15.01.2022 | Swab sampling: Oropharyngeal & Cloacal Swabs | **NEGATIVE**: Adélie penguin (100) |

| **Laboratory analyses** | | | |
| --- | --- | --- | --- |
| Investigators | Laboratory | Method | Protocol |
| Craig Cary,  Ian McDonald,  Roanna Richards-Babbage | University of Waikato | qPCR | Fresh guano was collected using the Thermofisher M4RT polyester swabs with 3 mLs of preservation solution and were frozen once collected.  Samples were thawed, shaken for 3 mins to thoroughly mix/breakup biomass into the solution. 1 mL of supernatant was taken and spun at 5,000 g for 30 secs to 10 mins depending on the sample viscosity.  50 uL of the supernatant was then extracted per sample using the MagMax^TM^ Viral/Pathogen Nucleic acid isolation kit (Thermofisher) in the presence of 2 µL of the 10,000 copies/µL Xeno™ RNA Control. A non-sample control of 2 µL of the 10,000 copies/ µL Xeno™ RNA Control in the presence of 50 uL of sterile PBS was included in each extraction batch as a positive extraction control. Samples were eluted in 20-25 µL of the kit supplied elution solution, and stored at -80 ^o^C until qPCR.  12.5 µL QPCR reactions were carried out using the Avian Influenza Virus RNA test kit (Thermofisher). 4 µL of each RNA sample was tested in duplicate. A standard curve was run in each assay for run to run comparison and to assess for low concentration of the target. Ct values were compared for the Xeno RNA extraction control and the AIV control with those of samples to determine the likelihood of inhibition and relative presence of target observed. |

### 6. Cape Bird

Location: 166.44 East, 77.44 South

**Breeding Season 2023/24**

| **Fieldwork** | | | |
| --- | --- | --- | --- |
| Investigators | Period | Type | Species (sample size) |
| Grant Ballard,  Annie E. Schmidt,  Jean Pennycook  Acknowledgement:  Christina Burnham, Nadia Swanson, Suzanne Winquist | 30.11.2023 – 01.12.2023 | Systematic observations | **NEGATIVE**: Adélie penguin, South polar skua (entire breeding colony) |

**Breeding Season 2022/23**

| **Fieldwork** | | | |
| --- | --- | --- | --- |
| Investigators | Period | Type | Species (sample size) |
| Arvind Varsani,  Jean Pennycook, Megan Elrod,  Dennis Jongomjit,  Amy Li | 22.11.2022 – 28.11.2022 | Observations | **NEGATIVE**: Adélie penguin, South polar skua (entire breeding colony) |

### 7. Cape Crozier

Location: 169.28 East, 77.46 South

**Breeding Season 2022/23**

| **Fieldwork** | | | |
| --- | --- | --- | --- |
| Investigators | Period | Type | Species (sample size) |
| Grant Ballard,  Annie E. Schmidt, Christina Burnham, Nadia Swanson | 11.07.2023 – 29.12.2023 | Systematic observations | **NEGATIVE**: Adélie penguin, Emperor penguin, South polar skua (entire breeding colony) |

**Breeding Season 2022/23**

| **Fieldwork** | | | |
| --- | --- | --- | --- |
| Investigators | Period | Type | Species (sample size) |
| Megan Elrod,  Grant Ballard,  Annie E. Schmidt,  Dennis Jongsomjit,  Amy Lee,  Aidan Cox | 12.11.2022 – 03.02.2023 | Observations | **NEGATIVE**: Adélie penguin, Emperor penguin, South polar skua (entire breeding colony) |

### 8. Cape Hallet, Victoria Land

Location: 170.23 East, 72.31 South

**Breeding Season 2022/23**

| **Fieldwork** | | | |
| --- | --- | --- | --- |
| Investigators | Period | Type | Species (sample size) |
| Craig Cary | 28/11/22 | Swab sampling of guano, observations | **NEGATIVE**: Adélie penguin (15) |

| **Laboratory analyses** | | | |
| --- | --- | --- | --- |
| Investigators | Laboratory | Method | Protocol |
| Craig Cary,  Ian McDonald,  Roanna Richards-Babbage | University of Waikato | qPCR | Fresh guano was collected using the Thermofisher M4RT polyester swabs with 3 mLs of preservation solution and were frozen once collected.  Samples were thawed, shaken for 3 mins to thoroughly mix/breakup biomass into the solution. 1 mL of supernatant was taken and spun at 5,000 g for 30 secs to 10 mins depending on the sample viscosity.  50 uL of the supernatant was then extracted per sample using the MagMax^TM^ Viral/Pathogen Nucleic acid isolation kit (Thermofisher) in the presence of 2 µL of the 10,000 copies/µL Xeno™ RNA Control. A non-sample control of 2 µL of the 10,000 copies/ µL Xeno™ RNA Control in the presence of 50 uL of sterile PBS was included in each extraction batch as a positive extraction control. Samples were eluted in 20-25 µL of the kit supplied elution solution, and stored at -80 ^o^C until qPCR.  12.5 µL QPCR reactions were carried out using the Avian Influenza Virus RNA test kit (Thermofisher). 4 µL of each RNA sample was tested in duplicate. A standard curve was run in each assay for run to run comparison and to assess for low concentration of the target. Ct values were compared for the Xeno RNA extraction control and the AIV control with those of samples to determine the likelihood of inhibition and relative presence of target observed. |

### 9. Damoy Point

Location: 63.51 West, 64.82 South

**Breeding Season 2023/24**

| **Fieldwork** | | | |
| --- | --- | --- | --- |
| Investigators | Period | Type | Species (sample size) |
| Grant Ballard | 03.02.2024 | Systematic observations | **NEGATIVE:** Gentoo penguin (most of breeding colony) |
| Megan Dewar,  Arvind Varsani | 25.02.2024 | Systematic observations | **NEGATIVE:** Chinstrap penguin, Brown skua |

### 10. Pléneau Island

Location: 64.05 West, 65.10 South

**Breeding Season 2023/24**

| **Fieldwork** | | | |
| --- | --- | --- | --- |
| Investigators | Period | Type | Species (sample size) |
| Grant Ballard | 03.02.2024 | Systematic observations | **NEGATIVE:** Gentoo penguin (most of breeding colony) |

### 11. Half Moon Island

Location: 59.90 West, 62.59 South

**Breeding Season 2023/24**

| **Fieldwork** | | | |
| --- | --- | --- | --- |
| Investigators | Period | Type | Species (sample size) |
| Grant Ballard | 05.02.2024 | Systematic observations | **NEGATIVE:** Chinstrap penguin (most of breeding colony), Gentoo penguin (15 molting) |

### 12. Neko Harbour

Location: 62.53 West, 64.85 South

**Breeding Season 2023/24**

| **Fieldwork** | | | |
| --- | --- | --- | --- |
| Investigators | Period | Type | Species (sample size) |
| Grant Ballard | 04.02.2024 | Systematic observations | **NEGATIVE:** Gentoo penguin (most of breeding colony) |

### 13. Cape Royds

Location: 169.28 East, 77.46 South

**Breeding Season 2023/24**

| **Fieldwork** | | | |
| --- | --- | --- | --- |
| Investigators | Period | Type | Species (sample size) |
| Jean Pennycook,  Grant Ballard,  Annie E. Schmidt, Suzanne Winquist | 09.11.2023 – 03.01.2024, | Systematic observations | **NEGATIVE:** Adélie penguin, South polar skua (entire breeding colony) |

**Breeding Season 2022/23**

| **Fieldwork** | | | |
| --- | --- | --- | --- |
| Investigators | Period | Type | Species (sample size) |
| Arvind Varsani,  Jean Pennycook,  David Ainley,  Grant Ballard,  Annie E. Schmidt,  Amelie Lescroel,  Aidan Cox,  Megan Elrod | 10.11.2022 – 14.01.2023, 06.02.2023 | Observations | **NEGATIVE**: Adélie penguin, South polar skua (entire breeding colony) |

### 14. Heronia Island

Location: 54.67 West, 63.43 South

**Breeding Season 2023/24**

| **Fieldwork** | | | |
| --- | --- | --- | --- |
| Investigators | Period | Type | Species (sample size) |
| Megan Dewar,  Arvind Varsani | 31.12.2023 | Swab samples | **NEGATIVE:** 6 dead Brown skuas |
| Megan Dewar,  Arvind Varsani | 31.12.2023 | Systematic observations | **NEGATIVE**: Adelie penguin (part of the 300000 pairs). Browns kua (~20), Giant petrel, Sheathbills |

| **Laboratory analyses** | | | |
| --- | --- | --- | --- |
| Investigators | Laboratory | Method | Protocol |
| Zoe Fowler  Arvind Varsani,  Meagan Dewar | KEMH Pathology and Food, Water & Environmental Laboratory  St Mary’s Walk, Stanley, Falkland Islands, FIQQ 1ZZ | qPCR | See laboratory protocol for Falkland Islands (same Laboratory and same protocol). |

### 15. Plaulet Island

Location: 55.78 West, 63.58 South

**Breeding Season 2023/24**

| **Fieldwork** | | | |
| --- | --- | --- | --- |
| Investigators | Period | Type | Species (sample size) |
| Megan Dewar,  Arvind Varsani | 31.12.2023 | Systematic observations | **NEGATIVE:** Adelie penguin, Brown skua (~20), Giant petrel, Sheathbill |

### 16. Devil Island

Location: 57.30 West, 63.53 South

**Breeding Season 2023/24**

| **Fieldwork** | | | |
| --- | --- | --- | --- |
| Investigators | Period | Type | Species (sample size) |
| Megan Dewar,  Arvind Varsani | 02.01.2023 | Systematic observations | **NEGATIVE:** Adelie penguin, Brown skua (~20) |

### 17. Cuverville Island

Location: 62.37 West, 64.68 South

**Breeding Season 2023/24**

| **Fieldwork** | | | |
| --- | --- | --- | --- |
| Investigators | Period | Type | Species (sample size) |
| Megan Dewar,  Arvind Varsani | 05.01.2023 | Systematic observations | **NEGATIVE:** Gentoo penguin, Brown skua (~20) |

### 18. Brown Base

Location: 62.87 West, 64.88 South

**Breeding Season 2023/24**

| **Fieldwork** | | | |
| --- | --- | --- | --- |
| Investigators | Period | Type | Species (sample size) |
| Megan Dewar,  Arvind Varsani | 06.01.2023 | Systematic observations | **NEGATIVE:** Gentoo penguin, Brown skua (~20), Antarctic shag (~5) |

### 19. Danco Island

Location: 62.60 West, 64.73 South

**Breeding Season 2023/24**

| **Fieldwork** | | | |
| --- | --- | --- | --- |
| Investigators | Period | Type | Species (sample size) |
| Megan Dewar,  Arvind Varsani | 06.01.2023  24.02.2024 | Systematic observations | **NEGATIVE:** Adelie penguin, Brown skua (~10) |

### 20. Brown Bluff

Location: 56.90 West, 63.52 South

**Breeding Season 2023/24**

| **Fieldwork** | | | |
| --- | --- | --- | --- |
| Investigators | Period | Type | Species (sample size) |
| Megan Dewar,  Arvind Varsani | 22.02.2023 | Systematic observations | **NEGATIVE:** Adelie penguin, Brown skua |

### 21. Orne Harbour

Location: 62.55 West, 64.63 South

**Breeding Season 2023/24**

| **Fieldwork** | | | |
| --- | --- | --- | --- |
| Investigators | Period | Type | Species (sample size) |
| Megan Dewar,  Arvind Varsani | 24.02.2023 | Systematic observations | **NEGATIVE:** Chinstrap penguin, Brown skua |

### 22. Cierva Cove

Location: 60.88 West, 64.15 South

**Breeding Season 2022/23**

| **Fieldwork** | | | |
| --- | --- | --- | --- |
| Investigators | Period | Type | Species (sample size) |
| Virginia Morandini  Josabel Belliure | 10.1.23  29.1.23 | Observation | **NEGATIVE**: Gentoo penguin, entire breeding colony |

### 23. Deception Island

Location: 62.96 West, 60.56 South

**Breeding Season 2022/23**

| **Fieldwork** | | | |
| --- | --- | --- | --- |
| Investigators | Period | Type | Species (sample size) |
| Virginia Morandini  Josabel Belliure | 24.12.23  14.2.23 | Observation | **NEGATIVE**: Chinstrap penguin, Vapor Col colony. Entire breeding colony (17.000 breeding pairs) |

### 24. Esperanza Base

Location: 56.99 West, 63.39 South

**Breeding Season 2022/23**

| **Fieldwork** | | | |
| --- | --- | --- | --- |
| Investigators | Period | Type | Species (sample size) |
| Virginia Morandini  Josabel Belliure | 16.1.23 | Observation | **NEGATIVE**: Adélie and gentoo penguin, entire breeding colonies |

### 25. Hannah Point, Livingston Island, South Shetland Islands

Location: 60.61 West, 62.56 South

**Breeding Season 2022/23**

| **Fieldwork** | | | |
| --- | --- | --- | --- |
| Investigators | Period | Type | Species (sample size) |
| Virginia Morandini  Josabel Belliure | 7.1.23  8.1.23 | Observation | **NEGATIVE**: Gentoo penguin and Adélie penguin. Entire breeding colonies |

### 26. Kerguelen

Location: 70.22 East, 49.35 South

**Breeding Season 2023/24**

| **Fieldwork** | | | |
| --- | --- | --- | --- |
| Investigators | Period | Type | Species (sample size) |
| Tierry Boulinier,  Jérémy Tornos,  Tristan Bralet | 20/11/2023-15/12/2023 | Systematic observation | **NEGATIVE**: Kelp gull (20), Brown skua (10), Northern giant petrel (10), Kerguelen hags (10), gentoo pengins (30). |
| Tierry Boulinier  Célia Lesage | 20/11/2023-01/03/202 | Opportunistic observation | **NEGATIVE**: Entire seabird breeding community, Péninsule Courbet, Golfe du Morbihan, Péninsule Jeanne d’Arc |

**Breeding Season 2022/23**

| **Fieldwork** | | | |
| --- | --- | --- | --- |
| Investigators | Period | Type | Species (sample size) |
| Jérémy Tornos,  Mathilde Lejeune | 24.12.2022-25/12/2023 | Systematic observation | **NEGATIVE**: Kelp gull (10), Brown skua (3), Northern giant petrel (5) |
| Tierry Boulinier  Jérémy Tornos,  Mathilde Lejeune | 01.09.2022-01.05.2022 | Opportunistic observation | **NEGATIVE**: Entire seabird breeding community, Péninsule Courbet, Golfe du Morbihan, Péninsule Jeanne d’Arc |

### 27. Marion Island, Prince Edward Islands

Location: 37.84 East, 46.88 South

**Breeding Season 2023/24**

| **Fieldwork** | | | |
| --- | --- | --- | --- |
| Investigators | Period | Type | Species (sample size) |
| Maelle Connan  Acknowledgement:  Michelle Risi,  Christopher Jones | Continuously (01.05.2023-29.02.2024) | Systematic observation | **NEGATIVE**: Wandering albatross, Grey-headed albatross, Northern giant petrels, and opportunistic observations on Brown skua, Kelp gull, Black-faced sheathbill |

**Breeding Season 2022/23**

| **Fieldwork** | | | |
| --- | --- | --- | --- |
| Investigators | Period | Type | Species (sample size) |
| Maelle Connan,  Lucy Smyth | Continuously (01.05.2022-30.04.2023) | Systematic observation | **NEGATIVE**: Wandering albatross, Grey-headed albatross, Northern and Southern giant petrels, Brown skua, Kelp gull, Black-faced sheathbill |

### 28. Penguin Island, South Shetland Islands

Location: 57.92 West, 62.10 South

**Breeding Season 2022/23**

| **Fieldwork** | | | |
| --- | --- | --- | --- |
| Investigators | Period | Type | Species (sample size) |
| Virginia Morandini  Josabel Belliure | 13.1.23 | Observation | **NEGATIVE:** Chinstrap penguin. Entire breeding colonies |

### 29. Possession Island, Crozet archipelago

Location: 51.86 East, 46.43 South

**Breeding Season 2022/23**

| **Fieldwork** | | | |
| --- | --- | --- | --- |
| Investigators | Period | Type | Species (sample size) |
| Tierry Boulinier,  Tristan Bralet | 06/11/2023 -06/12/2023 | Systematic observation | **NEGATIVE:** Brown skua (40), Northern giant petrel (30), Southern giant petrel (10), King penguin (80), Macaroni penguin (60), Southern rock hopper penguin (20), White-chinned petrel (20), Lesser sheathbill (90). |
| Tierry Boulinier,  Tristan Bralet | 06/11/2023-06/12/2023 | Observation | **NEGATIVE**: Entire seabird breeding community, in particular at Baie du Marin, Baie américaine, Pointe basse |

**Breeding Season 2022/23**

| **Fieldwork** | | | |
| --- | --- | --- | --- |
| Investigators | Period | Type | Species (sample size) |
| Jérémy Tornos,  Mathilde Lejeune | 25.08.2022-21-12-2022 | Systematic observation | **NEGATIVE:** Brown skua, Northern giant petrel, Southern giant petrel, King penguin, Macaroni penguin, Southern rock hopper penguin, White-chinned petrel, Lesser sheathbill |
| Tierry Boulinier  Jérémy Tornos,  Mathilde Lejeune | 01.09.2022-01.05.2022 | Observation | **NEGATIVE**: Entire seabird breeding community, in particular at Baie du Marin, Baie américaine, Pointe basse |

### 30. Stranger Point, King George Island, South Shetland Islands

Location: 58.26 West, 62.26 South

**Breeding Season 2022/23**

| **Fieldwork** | | | |
| --- | --- | --- | --- |
| Investigators | Period | Type | Species (sample size) |
| Virginia Morandini  Josabel Belliure | 14.1.23 | Observation | **NEGATIVE**: Gentoo penguin and Adélie penguin. Entire breeding colonies |

### 31. Ronge Island

Location: 62.68 West, 64.71 South

**Breeding Season 2022/23**

| **Fieldwork** | | | |
| --- | --- | --- | --- |
| Investigators | Period | Type | Species (sample size) |
| Virginia Morandini  Josabel Belliure | 18.1.23 | Observation | **NEGATIVE**: Gentoo penguin and Chinstrap penguin. Entire breeding colonies |

### 32. Yalour Island

Location: 64.16 West, 65.23 South

**Breeding Season 2022/23**

| **Fieldwork** | | | |
| --- | --- | --- | --- |
| Investigators | Period | Type | Species (sample size) |
| Virginia Morandini  Josabel Belliure | 28.1.23 | Observation | **NEGATIVE**: Adélie penguin. Entire breeding colonies |
